# Single-shot impulsive stimulated Brillouin microscopy by tailored ultrashort pulses

**DOI:** 10.1101/2024.08.07.606954

**Authors:** David Krause, John Boehm, Leon Liebig, Nektarios Koukourakis, Juergen W. Czarske

## Abstract

Brillouin microscopy has become an important tool for investigating the mechanical properties of tissue. The recently developed Impulsive stimulated Brillouin Scattering (ISBS) promises a label-free, non-invasive measurements of viscoelastic properties of transparent samples and offers the potential for a high temporal resolution. However, the spatial resolution of ISBS is currently limited, which hinders its transfer to real-world applications. Increasing the spatial resolution of ISBS leads to an increase in the energy density of the pump beams, which requires a balancing of the excitation parameters to stay below the phototoxic threshold. This paper focuses on the influences of different excitation parameters on the spatial, temporal and spectral resolution and their optimal values. Combined with the adoption of a noise suppressing window function, a measurement rate of 20µs/pixel in hydrogel is achieved, which is promising for fast 3D imaging. The presented advanced impulsive stimulated Brillouin microscopy can be applied for fast tissue elastography to-ward disease studies.

## Introduction

The study of mechanical properties of biological material provides important insights into its biological function (1, 2) and can characterize the pathological states of tissue (3). Consequently, mechanical mapping of tissue by medical imaging is a rapidly developing field that has emerged as a valuable tool in biomedical research and is already used in clinical practice as organlevel diagnostic assistance tool for various diseases such as prostate cancer (4, 5), liver fibrosis (6) and breast lesions (7). Many techniques have been developed, such as ultrasound elastography (8) and atomic force microscopy (AFM) (9). However, the acquisition of mechanical properties at sub-cellular resolution has long posed significant challenges, primarily due to the reliance of traditional techniques on direct physical contact or their insufficient cellular resolution. Optical elastography on the other hand offer various advantages over classical methods, being non-invasive, contactless, label-free, and can provide cellular or even sub-cellular resolution (3).

An emerging optical elastography technique is Brillouin microscopy (BM) which is based on the interaction of light (photons) with an acoustic wave (phonons) (10–14) and has become a highly appreciated technique for studying condensed matter systems (15–17). The spontaneous BM (SpBM) offers high spatial resolution, but is based on the interaction of photons with thermal noise which has a low power spectral density. This leads to long integration times and therefore SpBM making the application in clinics difficult. Despite these challenges line measurements at acquisition times in the range of 100-200 ms, which effectively equates to 1 ms/pixel, have been accomplished (18, 19).

The active stimulation of phonons, instead of the resonant spontaneous scattering of thermally generated phonons, can lead to an efficiency that is orders of magnitude higher and thus to faster imaging. Stimulated Brillouin scattering (SBS) utilizes two slightly detuned continuous-wave (cw) lasers for phonon excitation (pump and probe technique) (20–24) and achieves scanning times of ∼20 ms/pixel (22), but has limitations because it still requires the scanning of one excitation wavelength. Impulsive stimulated Brillouin scattering (ISBS) holds promise for achieving measurement rates in the MHz range, owing to the signal generation through the interaction of ultra-short laser pulses (25–30). The higher temporal resolution of ISBS, facilitated by the interaction of two pump pulses and a probe beam, can significantly reduce measurement duration’s. This advancement is crucial for both rapid imaging in clinical applications and fundamental research, such as studying morphogenesis at the organism level during gastrulation in *Drosophila* (19).

The interference fringes formed by two intersecting laser pulses create phonons at the intersection point, where the probe photons are inelastically scattered. The stiffness of the material is linked to the Doppler frequency shift of the inelastically scattered photons. A investigation was performed on the influences of the pulse and probe parameters on the signal-to-noise ratio (SNR) and it was concluded that the choice of high pulse energies offers the greatest leverage (31). However, when focusing sharper to increase the spatial resolution, the increasing excitation energy density needs to be taken into account to avoid phototoxic effects. Recently Li et al. (28) performed a similar study, where they investigated the influence of the probe beam parameters and acquisition time. In this paper, we expand the investigation on optimizing the ISBS excitation process, which requires balancing the influence of excitation parameters, like the repetition rate on the spatial and frequency resolution. The temporal resolution of ISBS is increased by incorporating the exponential window function (32) into the signal analysis process.

## Results

The primary objective of this work is to find a balance between multiple experimental parameters that optimize the SNR of ISBS, resulting in faster measurement rates while simultaneously maintaining a high spatial and spectral resolution. The novel approach of using a exponential window for the frequency analysis of Brillouin signals lowered the required number of averages per measurement and thereby increased the temporal resolution.

### A. Exponential window function

Spectral leakage is a fundamental problem of frequency analysis with the fast Fourier transform (FFT), where the power of one frequency leaks into its neighboring frequencies. This phenomenon is attributed to the underlying assumption of the discrete Fourier transform (DFT) that the acquired signal repeats itself infinitely, which can lead to abrupt transitions of the signal. Commonly a window functions is used, which weights each time point with a value defined by the window function and usually taper off to zero at the start and end of the signal, thereby minimizing spectral leakage. However the benefit of less leakage comes at the cost of a reduced frequency resolution. There exist a plethora of different windows, each optimal for different use cases, for instance, the Hann window is the optimal choice for two far apart frequency components with unequal power (33). In the context of impact testing signals, the exponential window is a widely utilized option, because it minimizes the attenuation at its strongest point (the beginning), while ensuring a gradual decline to zero at the end. These characteristics can help improve the SNR (32) and make the adoption of this window for the exponentially damped ISBS signal an ideal choice.

The exponential decay of the signal defines the linewidth of the resulting peak in the frequency spectrum, and by multiplying the signal with another exponential function, the linewidth is changed. Assume you have the following intensity signal

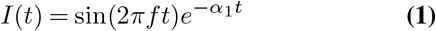

with *I*(*t*), *f* and *α*_1_ being the light intensity at the detector, frequency and the damping coefficient of the signal respectively. When this signal is then multiplied with the exponential window the resulting signal will be

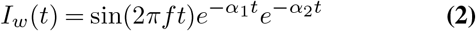

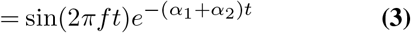

with *I*_*w*_(*t*), *α*_2_ being the windowed intensity signal and decay coefficient of the exponential window respectively. The linewidth of the peak in the frequency spectrum will now correspond to the sum of both decay coefficients 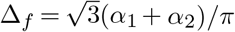(34). But as *α*_2_ is known a priori the calculation of the damping coefficient simply becomes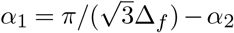.

To show the advantages of the exponential window in low SNR situations, simulations were carried out with different SNRs. The exponential window is compared with the results with a recently proposed technique for the frequency analysis based on adaptive noise suppression matrix pencils (ANMP), which promises to work well at low SNR levels (34). For each SNR level 100 signals are evaluated, allowing to estimate the variance of each method. The exponential window has the lowest variance both for the damping coefficient and the Brillouin frequency for low SNRs (Fig. 1 stays stable even for SNRs of -12 dB. At this SNR ANMP is not able to determine the linewidth, but achieves comparable variances compared to the exponential window at higher SNRs.

**Fig. 1.** The measured linewidth (top row) and peak frequency (bottom row) at different SNRs using the adaptive noise suppression matrix pencil (left column) or the exponential window (right column). The dashed line indicates the ground truth. The insets show the same plot with a zoomed in *y*-axis. For each SNR 100 signals with different noise are simulated.

Simply shortening the time signal to increase the SNR is not feasible, because under low SNR conditions the end of the Brillouin signal is difficult to determine. This makes the exponential window work especially good in situations with larger damping coefficients, where the signal decays quickly. To experimentally verify that the exponential window can increase the temporal resolution while accurately measuring the viscoelastic properties at low SNRs, measurements on ethanol were performed. A reference for the Brillouin frequency and linewdith of ethanol was created by averaging 10k measurements, which where taken with a pulse energy of 41 µJ. The averaged signal was evaluated with the FFT to obtain both the Brillouin frequency and the linewidth. Afterwards measurements at a reduced pulse energy with 20 averages were taken and the results of the analysis with ANMP and the exponential window are shown in Tab. 1. The ANMP method is capable of measuring the Brillouin frequency with an error margin of 10.53%, but it fails to accurately determine the signal’s linewidth. In contrast, the exponential window demonstrates the ability to measure both parameters, exhibiting a significantly lower error of 0.79% for the Brillouin frequency. This indicates that using the exponential window requires a SNR as low as -12.1 dB, thereby substantially reducing the required number of averages and optical power.

### B. Spatial resolution

The most straightforward approach to increase spatial resolution is to decrease the size of the focus by changing the focal lengths of the lens system. With the configuration depicted in Fig. 9 we studied an excitation cross-section diameter of 40 µm and a matched diameter for the probe beam. The fringe spacing in the excitation volume is 3 µm. The measurements were all conducted on a cuvette with 500 µm axial extent and lateral dimensions of 5 mm. We use signal strength, simply defined as the fitted peak height in the Fourier spectrum, as a measure.

As already derived in our previous paper (31), the signal strength dependence non linearly on the pulse energy and linearly on the probe power. The probe induced light power is limited due to phototoxicity, making the increase of the pulse energy more advantageous compared to the probe power. The reason for this lies in the fact, that for the same average power the pulsed excitation can have higher peak powers then with the continues probe beam. We at first choose a repetition rate of *k* = 1 kHz and averaged over *N* = 1000 samples. The signal strength increases non-linear with the pulse energy for all pulse lengths (Fig. 2 top left), making it advantageous to work with high energy pulses. However, starting from a certain peak power threshold we detect that for consecutive measurements the peak frequency started to fluctuate, which we attribute to phototoxicity. Thus decreasing the excitation volume size at constant pulse energies results in too high peak power densities. The peak power 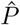 can be calculated by 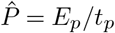, with *E*_*p*_ and *t*_*p*_ being the pulse energy and pulse length respectively. As plotted in the top right of Fig. 2, a good parameter choice for a high SNR is a long pulse length while keeping the pulse energy high to reduce the peak power. It should be noted, that all quantitative limits derived in this paper are dependent on the sample and on the sensitivity of the experimental setup.

**Fig. 2.**
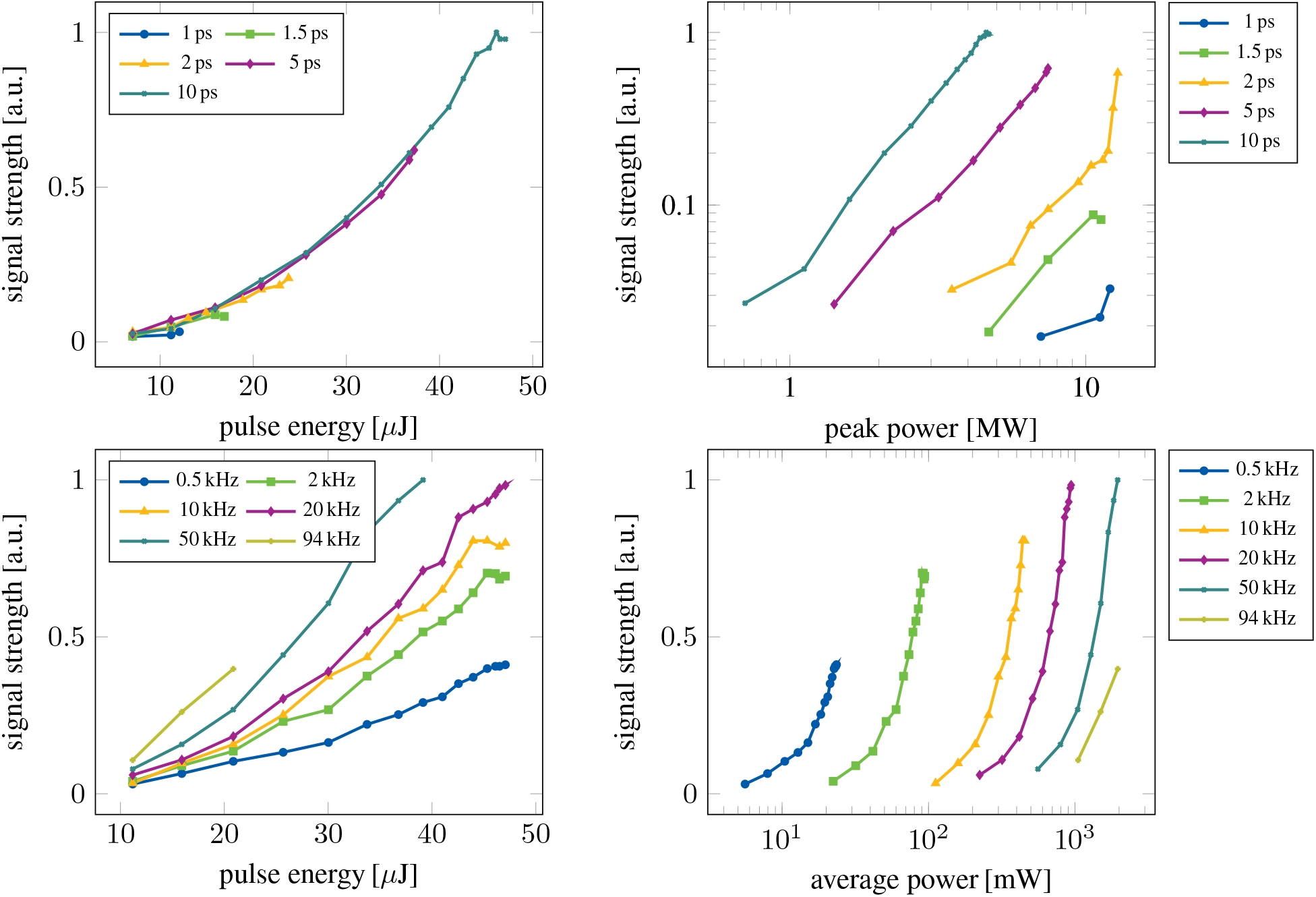
The dependence of the normalized signal strength on (top left) different pulse energies and pulse lengths, (top right) different peak powers and pulse lengths, (bottom left) different pulse energies and repetition rates and (bottom right) different average powers and repetition rates. For the top row the repetition rate was set to 1 kHz and in the bottom row the pulse length was kept constant at 10 ps for all measurements.

**Table 1.** Experimentally determined viscoelastic properties of ethanol. The values in parentheses are the relative error to the reference.

| Integration time (ms) | SNR (dB) | Spectral Analysis Method | Brillouin shift (MHz) | Linewidth (MHz) |
| --- | --- | --- | --- | --- |
| 1000 | 6 | Standard | 443.75 | 17.80 |
| 0.4 | -12.1 | Exponential window | 440.25 (0,79%) | 17.22 (3,26%) |
| 0.4 | -12.1 | ANMP | 492.06 (10,53%) | N/A |

Most importantly the behavior will be different for different pump wavelengths. In the subsequent analysis, we employ pulse duration’s of *t*_*p*_ = 10 ps, while changing both the repetition rate and the pulse energy to alter the average power 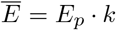. Both the pulse energy and the repetition rate increase the signal strength, illustrated in the bottom left in Fig. 2, which suggests picking the highest value for each possible.

But also for the average power a threshold is observed, above which the peak frequency started to fluctuate with an added drift of the peak frequency, which can be attributed to induction of heat. The repetition rate and pulse energy would be selected in such a way that the average power remains below the specified threshold, thus the determination of this threshold for a given sample is important. The trade-offs between choosing a high pulse energy or high repetition rate will be discussed in the next section.

Further decreasing the pump-beam diameter would increase the spatial resolution, but would elevate the danger of phototoxic effects in the sample.

### C. Temporal resolution

The achievable measurement rate is limited by the usable repetition rate and the required number of averages to create an adequate signal quality. For all aforementioned results, *N* = 1000 averages were used. Although the SNR increases with 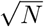 during averaging, the measurement time increases proportionally to *N*. Therefore, it is crucial to optimize the SNR of a single measurement to minimize the overall measurement time. Therefore, it is optimal to select the highest pulse energy possible and pick a repetition rate accordingly, which together stay below the threshold.

We found that single-shot measurements are possible at pulse energies of *E*_*p*_ = 35,µJ and a pulse repetition rate of 50 kHz with and without the exponential window. A relative precision of 0.3% is achieved, which we define here as adequate measurement quality, but the average power is very close to the maximum applicable dose. In order to not to impair the sample a 20 µJ pulse energy and 50 kHz repetition rate were chosen as parameters. Only when the single-shot measurement (Fig. 3a)) is evaluated with the exponential window (Fig. 3b)), it is possible to acquire the peak frequency and linewidth. There is still a peak present at the frequency of hydrogel in the spectrum when no window is used, but the SNR is too low for a reliable result.

**Fig. 3.**
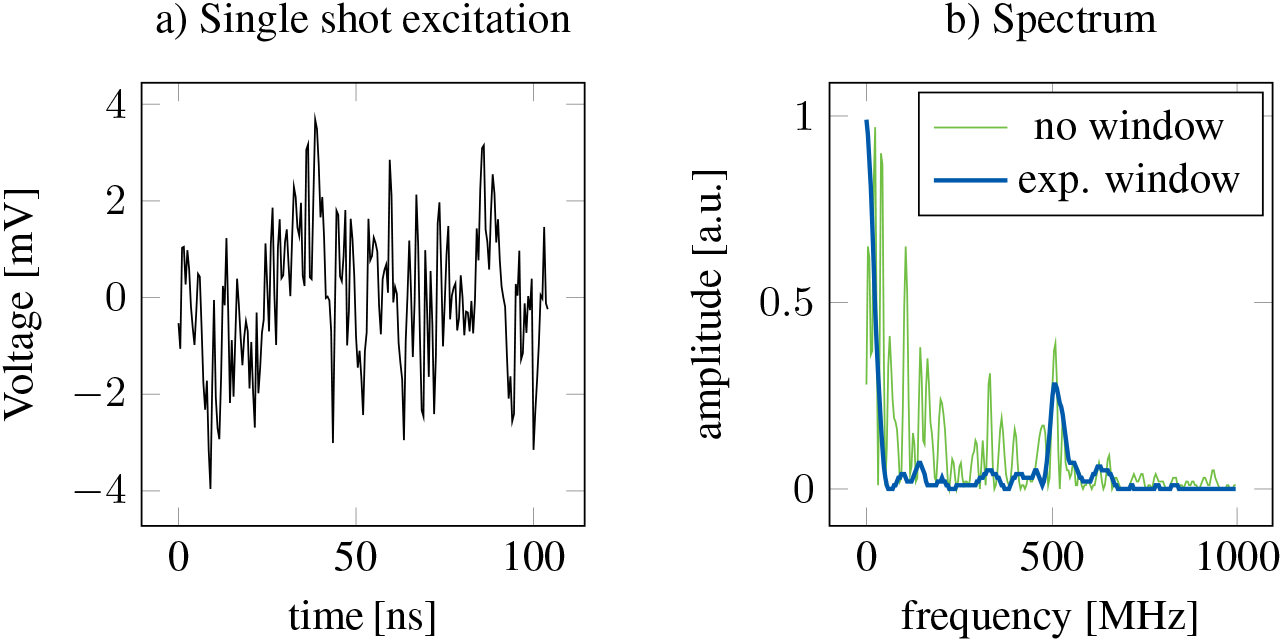
a) Time signal of a single-shot measurement and b) the corresponding frequency spectrum once evaluated without and with exponential window. c) The required number of averages to achieve a relative peak precision of 0.3 %, for different pulse energies and repetition rates.

We applied stage-scanning to record 2D images of a hydrogel-cube, freshly located in water with a coarse spatial resolution of 100 µm (Fig. 4). The hydrogel and the water frequencies can be clearly distinguished, see Fig. 4 on the right. With the measurement point rate of 50 kHz the theoretical frame rate for this 24 pixel image is 0.48 ms corresponding to a pixel measurement rate of 20 µs/pixel, which is around two orders of magnitudes faster then current then SpBM techniques.

**Fig. 4.**
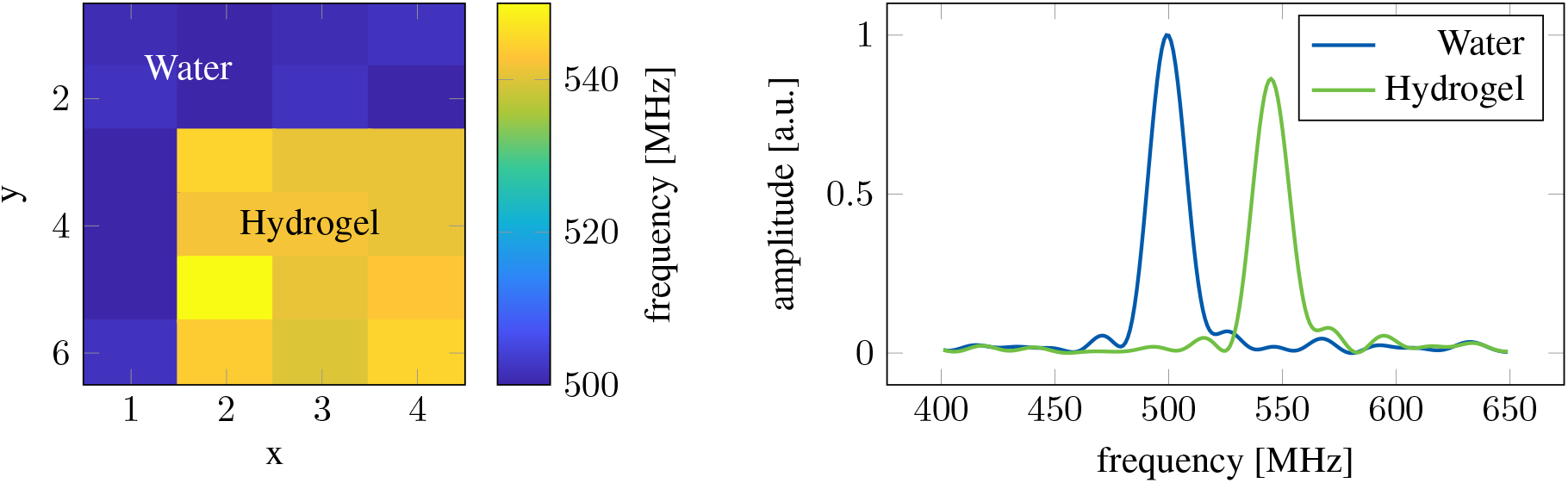
Image of a hydrogel cube embedded in water with the corresponding spectra on the right.

### D. Spectral resolution

The last subsections showed that ISBS has the potential to enable high-speed Brillouin measurements. However, due to the desired small measurement volume, the extent of the signal is also short in time, lasting approximately 40 ns in this example. According to the Cramer-Rao lower bound (CRLB), this short signal duration fundamentally limits the obtainable spectral resolution. Specifically, the CRLB indicates that the minimum variance of any unbiased estimator of the frequency is inversely proportional to the signal duration (35). The spectral resolution should not be confused with the resolution of the Fourier sampling, which can be enhanced by zero padding and improves the peak determination accuracy.

Hence, a small measurement volume is usable mainly in homogeneous media, where one frequency is present. If two materials are located in the measurement volume and the spectral resolution is not high enough to distinguish the corresponding two peaks, the measurement will be distorted by a shifted peak position, as the two peaks will merge to one peak with broader bandwidth. A high enough spectral resolution would enable resolving the two peaks. The only way to enhance the spectral resolution and to enable measurements of more complex samples is to increase the duration of the time signal, which can be achieved by sacrificing the resolution in *x*-direction(Fig. 5). Phonons across the whole cross section are excited and the ones created outside of the probe area propagate into the probing area and thereby extent the time signal. The extent of this effect strongly depends on the attenuation of the phonons and coupling efficiencies between the different media. The expansion in *x*-direction can be achieved by using a cylindrical lens to focus the pump beam on the grating. The expansion of the pump beam geometry does not only lead to higher spectral resolution as recently published in (36), but higher pulse energies can be used, as the expanded pump geometry spreads the deposited energy over a large area. The probe beam can keep its symmetric profile, which in our case has a full width at half maximum (FWHM) of approximately 40 µm (Fig. 5).

**Fig. 5.**
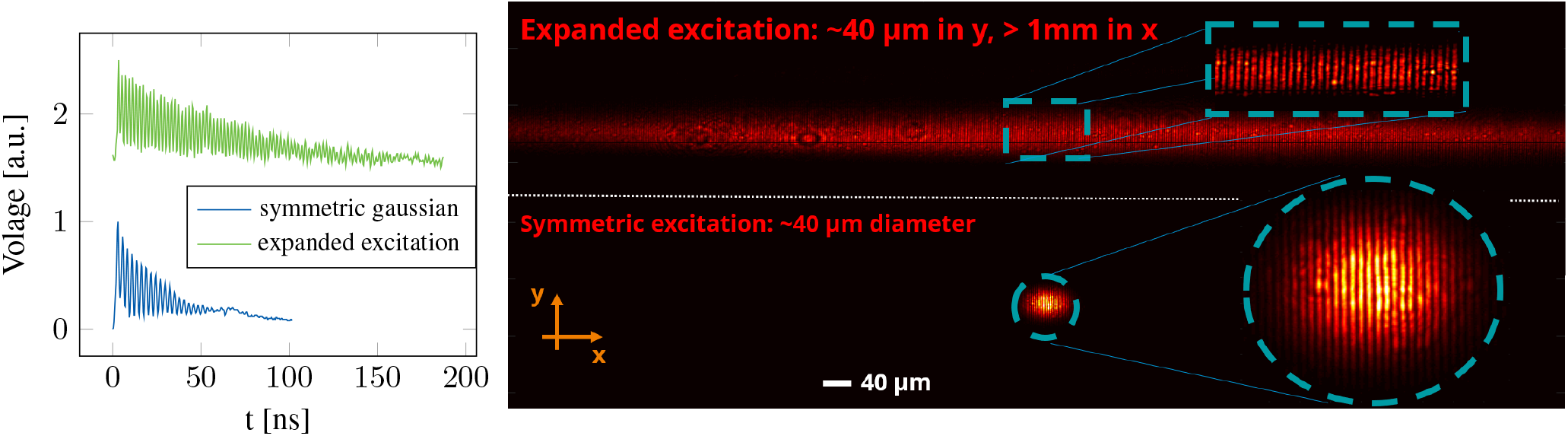
(left) Time signals for a symmetric gaussian and expanded excitation (offset added for clarity). (right) Images of the two excitation profiles.

If the excitation cross section overlaps at an interface of two different materials, it will create phonons in both materials with the same wavelength, because the same interference grating is imprinted in both materials. However, due to different sound velocities in the materials the phonons will have different frequencies. When phonons transmit through an interface they change their wavelength, but their frequency will stay constant. This frequency will be detected when the phonon passes through the probe area (Fig. 6a)). The detection of the frequency depends on the transmission efficiency of phonons through the interface and the attenuation of the phonons, rendering it therefore sample dependent. If the probe beam is in close proximity to the interface, the intersection between the probe and the interference grating under an angle also leads to the detection of both frequencies (Fig. 6b). The probe beam gets reflected from the grating periods in both materials. The strength of this volumetric effect depends on the intersection angle and the axial extent of the interference fringes.

**Fig. 6.**
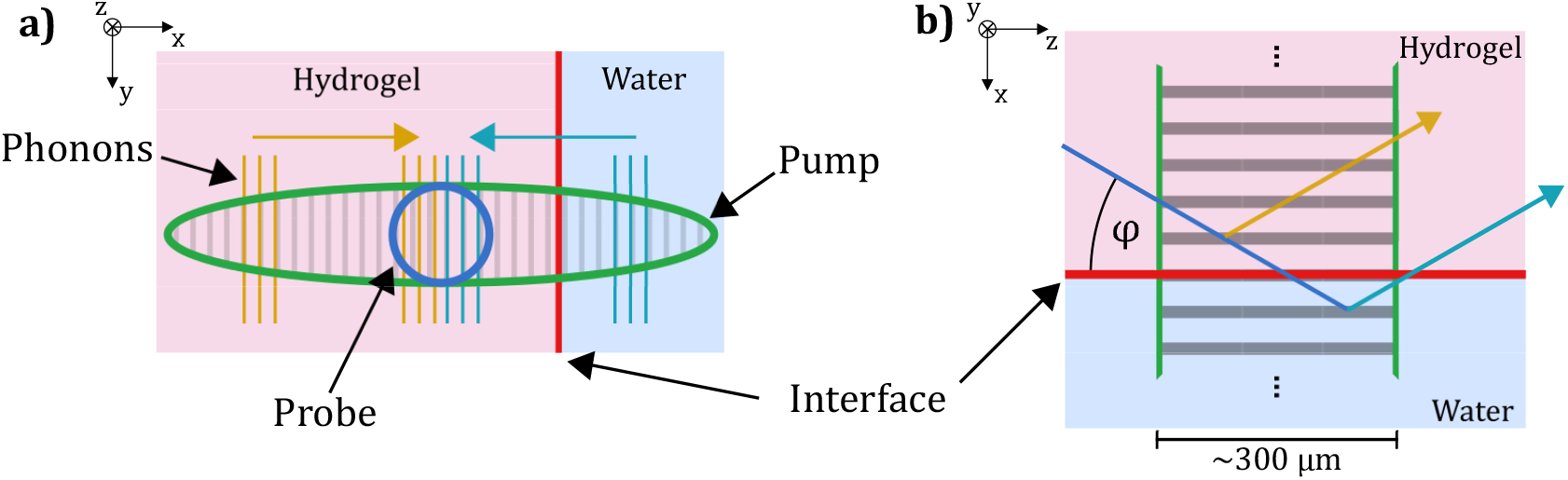
a) Schematic of an elliptical excitation in *x*-direction, which induces phonons in water and hydrogel, which after some time will reach the probe beam. When the in water created phonons transmit through the interface, they will propagate through the probe beam. b) Schematic of the volumetric effect, where due to the angle *φ* the probe beam is reflected both in water and hydrogel, even though the center of the probe beam is in hydrogel.

To verify the transmission of phonons, the measurement of a hydrogel cube in water was repeated, only with the line-shaped excitation profile. To study the behavior of the signals for different distances to the interface, we scan the sample laterally, thereby changing the relative position at which the probe beam enters the sample. The resulting spectra of the horizontal scan are shown in Fig. 7. Starting far from the interface, the ISBS signal shows only the frequency of hydrogel *f* = 500 MHz. Moving closer to the interface, the excited phonons in water transmit through the water/hydrogel interface and propagate through the probe area, leading to the detection of the water frequency (493 MHz). The position, where the two peaks are of equal height, corresponds to the interface position. In our case this would be at the position *x* = 90 µm. In the vicinity of the interface, both the volume effect and the crosstalk effect superimpose, whereas in the outer regions only the crosstalk effect exists. The data suggest that the crosstalk in that sample is limited to ∼90 µm around the interface, because at *x* = 190 µm only the water frequency is detected. This indicates a strong attenuation of the acoustic waves or a weak coupling between the two materials. The two different materials can be clearly distinguished after ∼180 µm, corresponding to the effective spatial resolution along the *x*-axis.

**Fig. 7.**
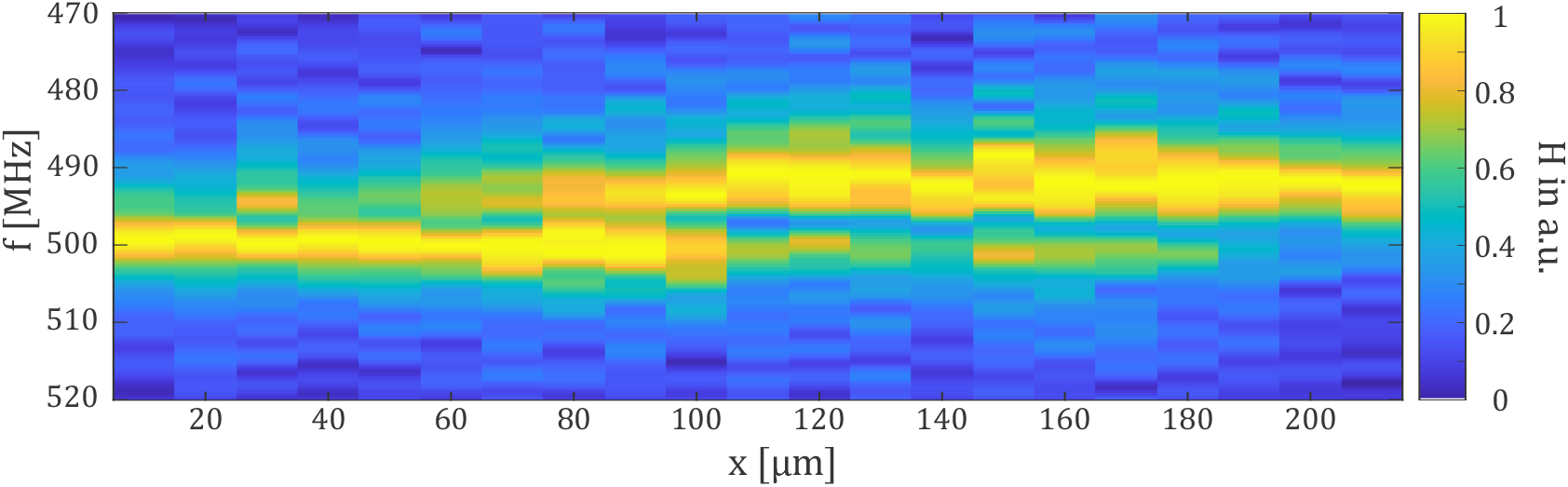
Normalized spectra at different *x* positions. The probe beam is scanned over an interface between hydrogel and water, starting in hydrogel. The amplitude *H* of each spectrum is normalized independently.

## Discussion

The noise suppression capability of the exponential window comes at the cost of a reduced frequency resolution of the system. The reduction in frequency resolution depends on the decay coefficient of the exponential and can be tuned, resulting in a frequency resolution which is up to 3 times larger compared to the analysis without a window. By tuning the decay coefficient you trade off a higher frequency resolution against the noise suppression capability and spectral leakage. Small decay coefficients offer a higher frequency resolution, but increased spectral leakage, while using larger decay coefficients improves noise suppression but at a lower frequency resolution. Thus, according to the SNR of the signal the decay coefficient should be picked. For low SNR signals a high decay coefficient is beneficial due to the noise suppression, but for higher SNR signals a lower decay coefficient can be viable.

Crucial parameters for measurements are the maximum allowable light dose and the sensitivity of the detection. These parameters limit the usable peak and average powers of the pump laser and thus the SNR and the achievable measurement rates. It is generally desirable to approach the imposed limits of the sample in order to obtain the best possible SNR. For long-time observations, the pulse energy should be reduced because at a repetition rate of 50 kHz and a pulse energy of 20 µJ the induced heating changes the speed of sound inside the sample.

For a fixed frequency resolution the speed of sound resolution could be further improved by decreasing the interference fringe distance *d*. This changes the expected frequency *f*_*e*_ following the equation for excitation *f*_*e*_ = *v*_*u*_*/d*, with *v*_*u*_ being the ultrasonic velocity. This is a linear equation which grows steeper with decreasing interference fringe spacing, see Fig. 8. Following the equation of the fringe spacing *d* = *λ*_pump_*/*(2 sin *θ*), with *θ* being half the intersection angle of the pump beams, *d* is at its smallest value when the angle between the pump beams is increased to 180°, leading to counter propagating pump beams. When two materials exhibit similar *v*_*u*_’s, it is easier to distinguish both materials with small fringe distances because a small change in *v*_*u*_ leads to a larger difference in frequency compared to bigger fringe distances, which in theory is easier to detect.

**Fig. 8.**
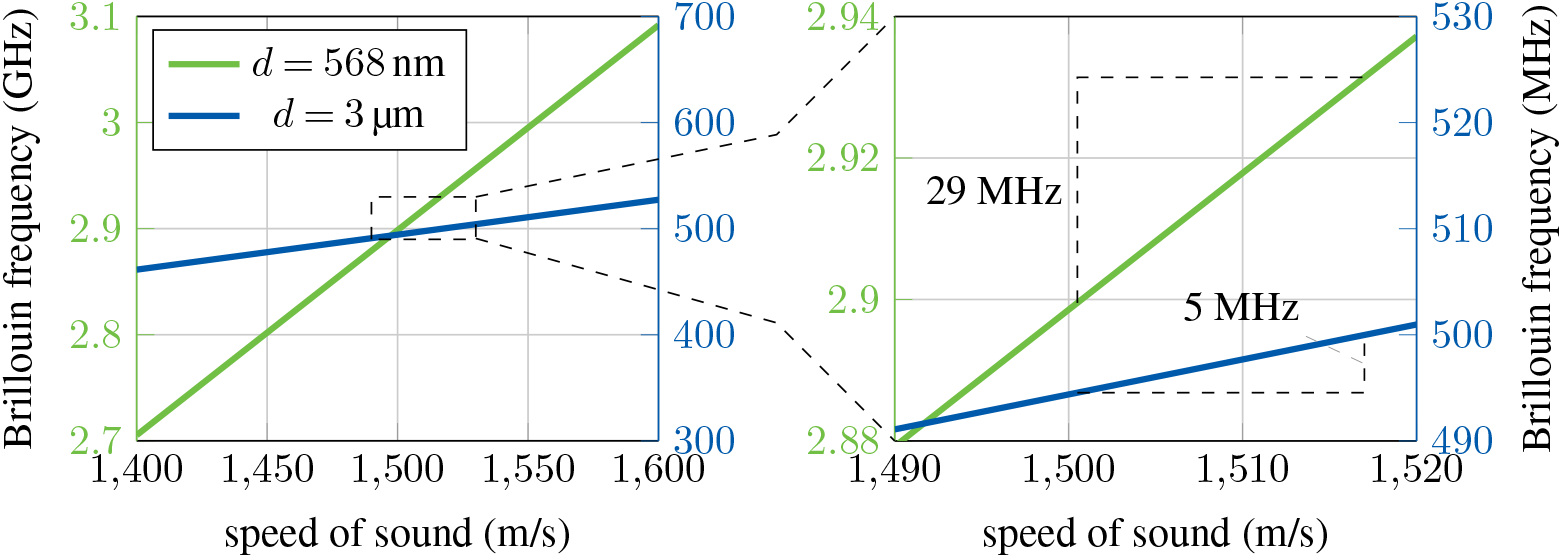
Relationship of the speed of sound to the Brillouin frequency for a grating period of *d* = 3 µm and *d* = 568 nm.

**Fig. 9.**
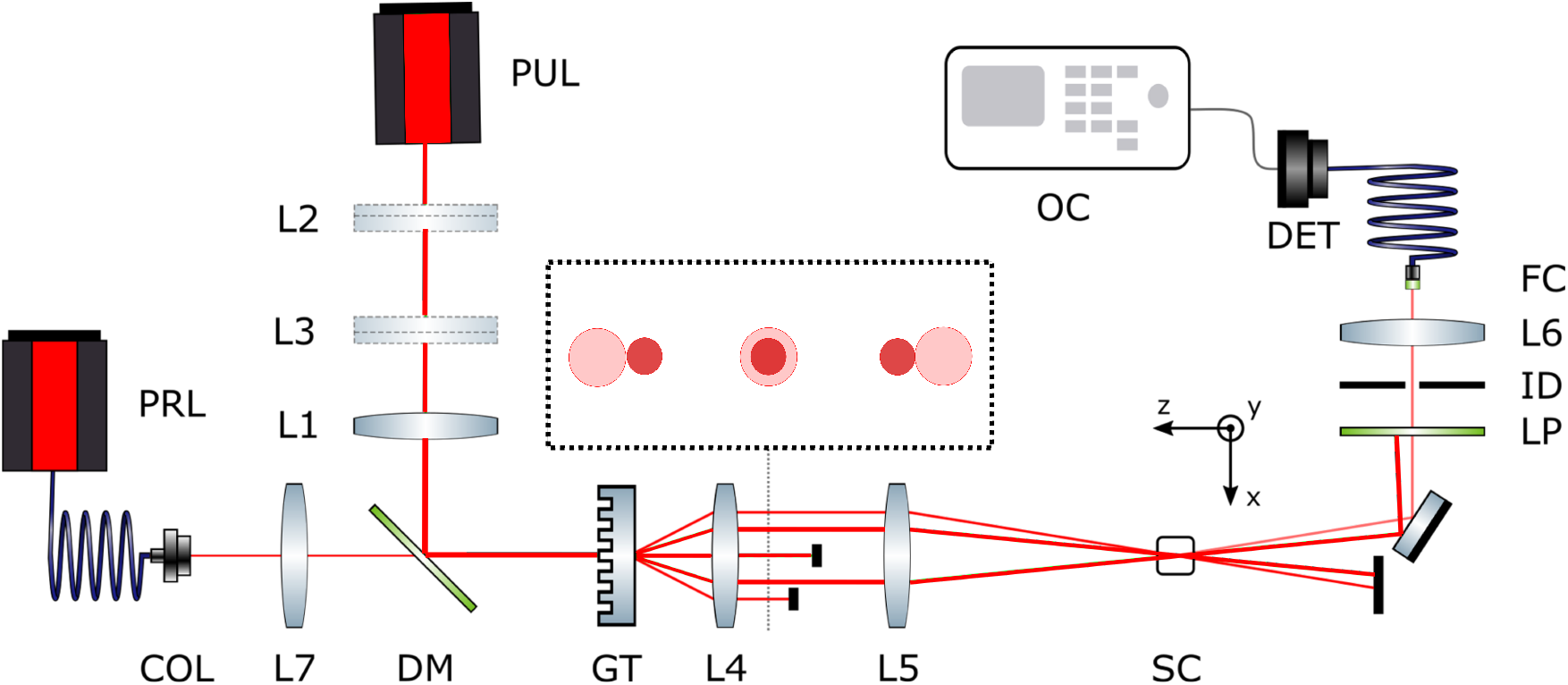
ISBS microscope setup: PUL excitation laser (1035 nm, Coherent Monaco); PRL probe laser (895 nm, DL100 Toptica Photonics); DM dichroic mirror (DMLP650 Thorlabs); COL collimator; L1 to L6 achromatic lenses (AC245-Series Thorlabs); GT grating (custom, grating constant 8 µm); SC sample container (Fused Quartz Cuvettes Thorlabs); ID iris diaphragm; LP long-pass (FEL0850 Thorlabs); DET detector (APD, Femto Berlin); OC oscilloscope.

The advantages of the steeper slope are two-fold regarding the spatial and speed of sound resolution respectively. Under the assumption that the phonon attenuation is approximately constant for the frequency range used, the signal duration for a given spatial resolution is constant. This means that the spectral resolution is also constant, resulting in a higher speed of sound resolution. There exists a relationship of the phonon frequency to the attenuation of phonons in solids (37), but it has to be investigated on how it affects the signal duration in liquid or biological samples. If on the other hand the improved speed of sound resolution is not required, the smaller fringe spacing has the benefit of requiring shorter time signals to achieve the same resolution in the speed of sound. In the case of a hydrogel cube embedded in water, which have a difference in the speed of sound of ∼17 m/s(1,1 %), the frequency difference between the two materials in the crossed beam setup would be 7 MHz and in the counter propagating setup would result in 29 MHz (Fig. 8 on the right). Measurement of such a difference would require a ∼140 ns long signal in the crossed beams setup, compared to only 35 ns in the counter propagating setup. This would allow for a higher spatial resolution, because the time signal shortens with decreasing focus size. For a given speed of sound resolution, the counter propagating setup thus enables a higher spatial resolution. But the approach with the counter propagating pump is harder to align than the setup depicted in Fig. 9, where the probe and pump beam automatically fulfill the Bragg condition if focused correctly at the grating.

When modifying the pump beam profile for a better spectral resolution and SNR, it is crucial to bear in mind the impact on the spatial resolution. Altering the initial pump profile by expanding it along the *x*-axis using a cylindrical lens leads to enhancements in spectral resolution and spatial resolution along the *y*-axis. However, it also results in a notable decrease in spatial resolution along the *x*-axis. This decline is mostly influenced by the crosstalk of the different media and strongly sample dependent. The transmission efficiency of the interface and the attenuation of the material affect the amount of crosstalk. In our case this leads to a spatial resolution of ∼180 µm in *x*-direction.

The final challenge to enable fast imaging is to find an alternative to stage scanning, which limits the frame rate at the moment. The hurdle is, that during scanning of the probe beam the Bragg condition needs to be fulfilled. Thus, the beams incident to the grating should just shift laterally orthogonal to the grating, which can be realized with a 2D galvo scanning system.

## Conclusion

In this paper we investigated the influences of the excitation parameters of ISBS on the SNR and how this effects the temporal and spatial resolution. High temporal resolution is important to achieve high-throughput measurements. The pulse energy and average power are the most significant excitation parameters for increasing the signal strength. Using longer pulses length lowers the peak power of the pulse, which increases the maximum allowed pulse energy. Through the introduction of an expanded pump profile in the *x*-direction the spectral resolution got improved by a factor of ∼4 with probe beam size of ∼40×40 µm^2^. The probe beam size is currently limited by the focal length of the used optics, but can be further improved to ∼14×14 µm^2^ with lenses with shorter focal lengths (28). By also spreading the pulse energy over a larger area, higher pulse energies are possible, which positively influence the SNR. The here presented temporal resolution of 20µs/pixel of ISBS is already two orders of magnitude faster compared to SpBS. In combination with a detector array, line measurements of 100 pixel would result in an effective measurement rate of 200 ns/pixel, which would enable real-time imaging. Therefore, the here presented work brings ISBS one step closer to real world applications.

## Materials and methods

ISBS microscopy relies on creating a transient density grating using an ultra-short pulsed laser and probing it through Bragg diffraction with a continuous wave (cw) laser. The pulsed laser beam is divided into two coherent beams via a diffraction grating. The interference of these coherent laser beams forms an interference fringe system (38), with fringe spacing *d* determined by the excitation beam’s wavelength *λ*_pump_ and the half-crossing angle *ϕ*_pump_:

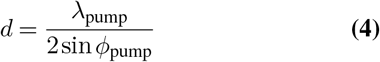

The impulsive excitation of a standing acoustic wave is induced by either electrostrictive or thermal coupling of the laser pulse to the material within the measurement volume. This density variation also alters the refractive index. Consequently, a portion of the probe beam is diffracted at a temporally varying refractive index grating under the Bragg condition, resulting in intensity modulation. For electrostrictive excitation, the expected modulation frequency is *f*_2_ = 2*c*_*S*_*/d*, where *c*_*S*_ is the sound velocity in the sample. If the signal is thermally excited, the expected frequency is *f*_1_ = *c*_*S*_*/d* (39). Electrostriction exerts a force on matter along the gradient of the absolute electric field strength (40), causing displacement of matter towards regions of high light intensity. Thermal excitation heats the matter in regions of higher light intensity, leading to impulsive expansion. The resulting temporal changes in matter are governed by thermodynamic material equations. Heterodyne detection can also be employed in ISBS microscopy (41). In this scenario, the probe beam reflected by the standing wave and another coherent part of the probe beam are superimposed on the photodetector. This additional part can be intentionally generated or arise inadvertently in the setup. For purely electrostrictive excitation with heterodyne detection, the expected frequency is *f*_1_. Both frequencies can be utilized to measure sound velocity and, consequently, deduce mechanical properties.

The experimental setup is depicted in Fig. 9. It is essentially the same architecture that was used in our previous work (31), with the only difference being that we now use pump laser with *λ*_pump_ = 1035 nm, while the probe beam is still at *λ*_probe_ = 895 nm (DL100 Toptica Photonics). The pump laser is a Coherent Monaco, which offers a tunable pulse-length, repetition-rate and pulse energy. The probe beam diffraction orders propagate between the diffraction orders of the grating for the pump laser. Both pump and probe beams are focused on the diffraction grating (GT) and the *±*1 diffraction orders are focused with the 4f-system to create the measurement volume. The choice of lenses is essential for the ISBS process. The −1 diffraction order of the probe beam is used to adjust the optical setup, but is later blocked to only detect the Brillouin scattered light. After the measurement volume, a long-pass filter removes the remaining pump light in the optical pathway and the Brillouin scattered probe beam is focused on an avalanche photodiode (APD). The oscilloscope (Tektronix MSO64B) reads out the time signal from the APD and sends it to a PC for analysis.

At this stage the signal contains a damped oscillation with an added decaying DC term. The exponential decaying DC term should be removed by fitting an exponential decaying DC term to the data, because it leads to a broad peak around the DC in the frequency spectrum, which makes the detection of Brillouin peaks difficult.

## ACKNOWLEDGEMENTS

The authors would like to thank DFG for their funding (CZ 55/44-1) and Anna Taubenberger (Biotechnology Center, TU Dresden) for providing hydrogel samples.

The authors declare no conflicts of interest.

## References

1. Brendan F. Kennedy, Philip Wijesinghe, and David D. Sampson. The emergence of optical elastography in biomedicine. Nature Photonics, 11(4):215–221, apr 2017. doi: 10.1038/nphoton.2017.6.

2. Nektarios Koukourakis, Felix Wagner, Stefan Rothe, Mike O Karl, and Jürgen W Czarske. Investigation of human organoid retina with digital holographic transmission matrix measurements. Light: Advanced Manufacturing, 3(2):211–225, 2022.

3. Nichaluk Leartprapun and Steven G. Adie. Recent advances in optical elastography and emerging opportunities in the basic sciences and translational medicine [invited]. Biomedical Optics Express, 14(1):208, dec 2022. doi: 10.1364/boe.468932.

4. J.-M. Correas, A.-M. Tissier, A. Khairoune, G. Khoury, D. Eiss, and O. Hélénon. Ultrasound elastography of the prostate: State of the art. Diagnostic and Interventional Imaging, 94(5):551–560, may 2013. doi: 10.1016/j.diii.2013.01.017.

5. Leo Pallwein, Fritz Aigner, Ralph Faschingbauer, Eva Pallwein, Germar Pinggera, Georg Bartsch, Georg Schaefer, Peter Struve, and Ferdinand Frauscher. Prostate cancer diagnosis: value of real-time elastography. Abdominal Imaging, 33(6):729–735, jan 2008. doi: 10.1007/s00261-007-9345-7.

6. Ounali S Jaffer, Phillip F C Lung, Diana Bosanac, Aarti Shah, and Paul S Sidhu. Is ultrasound elastography of the liver ready to replace biopsy? a critical review of the current techniques. Ultrasound, 20(1):24–32, jan 2012. doi: 10.1258/ult.2011.011043.

7. Ako Itoh, Ei Ueno, Eriko Tohno, Hiroshi Kamma, Hideto Takahashi, Tsuyoshi Shiina, Makoto Yamakawa, and Takeshi Matsumura. Breast disease: Clinical application of US elastography for diagnosis. Radiology, 239(2):341–350, may 2006. doi: 10.1148/radiol.2391041676.

8. Rosa MS Sigrist, Joy Liau, Ahmed El Kaffas, Maria Cristina Chammas, and Juergen K Willmann. Ultrasound elastography: review of techniques and clinical applications. Theranostics, 7(5):1303, 2017.

9. Helene OB Gautier, Amelia J Thompson, Sarra Achouri, David E Koser, Kathrin Holtzmann, Emad Moeendarbary, and Kristian Franze. Atomic force microscopy-based force measurements on animal cells and tissues. In Methods in cell biology, volume 125, pages 211–235. Elsevier, 2015.

10. Giuseppe Antonacci, Timon Beck, Alberto Bilenca, Jürgen Czarske, Kareem Elsayad, Jochen Guck, Kyoohyun Kim, Benedikt Krug, Francesca Palombo, Robert Prevedel, and Giuliano Scarcelli. Recent progress and current opinions in brillouin microscopy for life science applications. Biophysical Reviews, 12(3):615–624, may 2020. doi: 10.1007/s12551-020-00701-9.

11. Raimund Schlüßler, Stephanie Möllmert, Shada Abuhattum, Gheorghe Cojoc, Paul Müller, Kyoohyun Kim, Conrad Möckel, Conrad Zimmermann, Jürgen Czarske, and Jochen Guck. Mechanical mapping of spinal cord growth and repair in living zebrafish larvae by brillouin imaging. Biophysical journal, 115(5):911–923, 2018.

12. Jürgen Czarske and Harald Müller. Heterodyne detection technique using stimulated brillouin scattering and a multimode laser. Optics letters, 19(19):1589–1591, 1994.

13. Zachary N Coker, Maria Troyanova-Wood, Zachary A Steelman, Bennett L Ibey, Joel N Bixler, Marlan O Scully, and Vladislav V Yakovlev. Brillouin microscopy monitors rapid responses in subcellular compartments. PhotoniX, 5(1):9, 2024.

14. Conrad Moeckel, Timon Beck, Sara Kaliman, Shada Abuhattum, Kyoohyun Kim, Julia Kolb, Daniel Wehner, Vasily Zaburdaev, and Jochen Guck. Estimation of the mass density of biological matter from refractive index measurements. bioRxiv, pages 2023–12, 2023.

15. JG Dil. Brillouin scattering in condensed matter. Reports on Progress in Physics, 45(3):285, 1982.

16. Zijie Hua, Dexin Ba, Dengwang Zhou, Yijia Li, Yue Wang, Xiaoyi Bao, and Yongkang Dong. Non-destructive and distributed measurement of optical fiber diameter with nanometer resolution based on coherent forward stimulated brillouin scattering. Light: Advanced Manufacturing, 2(4):373–384, 2021.

17. Martina Alunni Cardinali, Silvia Caponi, Maurizio Mattarelli, and Daniele Fioretto. Brillouin scattering from biomedical samples: the challenge of heterogeneity. Journal of Physics: Photonics, 2024.

18. Jitao Zhang, Milos Nikolic, Kandice Tanner, and Giuliano Scarcelli. Rapid biomechanical imaging at low irradiation level via dual line-scanning brillouin microscopy. Nature methods, 20:677–681, 5 2023. ISSN 1548-7105. doi: 10.1038/s41592-023-01816-z.

19. Carlo Bevilacqua, Juan Manuel Gomez, Ulla-Maj Fiuza, Chii Jou Chan, Ling Wang, Sebastian Hambura, Manuel Eguren, Jan Ellenberg, Alba Diz-Muñoz, Maria Leptin, and Robert Prevedel. High-resolution line-scan brillouin microscopy for live imaging of mechanical properties during embryo development. Nature methods, 20:755–760, 5 2023. ISSN 1548-7105. doi: 10.1038/s41592-023-01822-1.

20. Itay Remer and Alberto Bilenca. Background-free brillouin spectroscopy in scattering media at 780 nm via stimulated brillouin scattering. Optics Letters, 41(5):926–929, 2016.

21. Itay Remer, Roni Shaashoua, Netta Shemesh, Anat Ben-Zvi, and Alberto Bilenca. High-sensitivity and high-specificity biomechanical imaging by stimulated brillouin scattering microscopy. Nature Methods, 17(9):913–916, 2020.

22. Fan Yang, Carlo Bevilacqua, Sebastian Hambura, Ana Neves, Anusha Gopalan, Koki Watanabe, Matt Govendir, Maria Bernabeu, Jan Ellenberg, Alba Diz-Muñoz, Simone Köhler, Georgia Rapti, Martin Jechlinger, and Robert Prevedel. Pulsed stimulated brillouin microscopy enables high-sensitivity mechanical imaging of live and fragile biological specimens. Nature Methods, pages 1–9, 10 2023. ISSN 1548-7091. doi: 10.1038/s41592-023-02054-z.

23. Charles W Ballmann, Jonathan V Thompson, Andrew J Traverso, Zhaokai Meng, Marlan O Scully, and Vladislav V Yakovlev. Stimulated brillouin scattering microscopic imaging. Scientific reports, 5(1):18139, 2015.

24. Tian Li, Fu Li, Xinghua Liu, Vladislav V Yakovlev, and Girish S Agarwal. Quantum-enhanced stimulated brillouin scattering spectroscopy and imaging. Optica, 9(8):959–964, 2022.

25. Charles W. Ballmann, Zhaokai Meng, Andrew J. Traverso, Marlan O. Scully, and Vladislav V. Yakovlev. Impulsive brillouin microscopy. Optica, 4:124, 1 2017. ISSN 2334-2536. doi: 10.1364/OPTICA.4.000124.

26. Benedikt Krug, Nektarios Koukourakis, and Juergen W Czarske. Impulsive stimulated brillouin microscopy for non-contact, fast mechanical investigations of hydrogels. Optics express, 27(19):26910–26923, 2019.

27. Taoran Le, Jiarui Li, Haoyun Wei, and Yan Li. Speed of sound measurement and mapping in transparent materials by impulsive stimulated brillouin microscopy. Journal of Physics: Photonics, 2024.

28. Jiarui Li, Taoran Le, Hongyuan Zhang, Haoyun Wei, and Yan Li. High-speed impulsive stimulated brillouin microscopy. Photonics Research, 12 2023.

29. Zhaokai Meng, Georgi I Petrov, and Vladislav V Yakovlev. Flow cytometry using brillouin imaging and sensing via time-resolved optical (bistro) measurements. Analyst, 140(21):7160–7164, 2015.

30. Charles W Ballmann, Zhaokai Meng, and Vladislav V Yakovlev. Nonlinear brillouin spectroscopy: what makes it a better tool for biological viscoelastic measurements. Biomedical optics express, 10(4):1750–1759, 2019.

31. Benedikt Krug, Nektarios Koukourakis, Jochen Guck, and Jürgen Czarske. Nonlinear microscopy using impulsive stimulated brillouin scattering for high-speed elastography. Optics express, 30:4748–4758, 2 2022. ISSN 1094-4087. doi: 10.1364/OE.449980.

32. W. Fladung and R. Rost. Application and correction of the exponential window for frequency response functions. Mechanical Systems and Signal Processing, 11:23–36, 1 1997. ISSN 08883270. doi: 10.1006/mssp.1996.0084.

33. KM Muraleedhara Prabhu. Window functions and their applications in signal processing. Taylor & Francis, 2014.

34. Jiarui Li, Hongyuan Zhang, Minjian Lu, Haoyun Wei, and Yan Li. Sensitive impulsive stimulated brillouin spectroscopy by an adaptive noise-suppression matrix pencil. Optics Express, 30:29598, 8 2022. ISSN 1094-4087. doi: 10.1364/OE.465106.

35. Jürgen W Czarske. Statistical frequency measuring error of the quadrature demodulation technique for noisy single-tone pulse signals. Measurement Science and Technology, 12 (5):597, 2001.

36. Sean P O’Connor, Dominik A Doktor, Marlan O Scully, and Vladislav V Yakovlev. Spectral resolution enhancement for impulsive stimulated brillouin spectroscopy by expanding pump beam geometry. Optics express, 31:14604–14616, 4 2023. ISSN 1094-4087. doi: 10.1364/OE.487131.

37. TC Zhu, Humphrey J Maris, and J Tauc. Attenuation of longitudinal-acoustic phonons in amorphous sio2 at frequencies up to 440 ghz. Physical Review B, 44(9):4281, 1991.

38. Philipp Günther, Robert Kuschmierz, Thorsten Pfister, and Jürgen Czarske. Distance measurement technique using tilted interference fringe systems and receiving optic matching. Optics letters, 37(22):4702–4704, 2012.

39. Stefan Schlamp, Hans G Hornung, Thomas H Sobota, and Eric B Cummings. Accuracy and uncertainty of single-shot, nonresonant laser-induced thermal acoustics. Applied Optics, 39 (30):5477–5481, 2000.

40. Robert W Boyd, Alexander L Gaeta, and Enno Giese. Nonlinear optics. In Springer Handbook of Atomic, Molecular, and Optical Physics, pages 1097–1110. Springer, 2008.

41. AA Maznev, KA Nelson, and JA Rogers. Optical heterodyne detection of laser-induced gratings. Optics letters, 23(16):1319–1321, 1998.

